# Understanding human amygdala function with artificial neural networks

**DOI:** 10.1101/2024.07.29.605621

**Authors:** Grace Jang, Philip A. Kragel

## Abstract

The amygdala is a cluster of subcortical nuclei that receives diverse sensory inputs and projects to the cortex, midbrain and other subcortical structures. Numerous accounts of amygdalar contributions to social and emotional behavior have been offered, yet an overarching description of amygdala function remains elusive. Here we adopt a computationally explicit framework that aims to develop a model of amygdala function based on the types of sensory inputs it receives, rather than individual constructs such as threat, arousal, or valence. Characterizing human fMRI signal acquired as participants viewed a full-length film, we developed encoding models that predict both patterns of amygdala activity and self-reported valence evoked by naturalistic images. We use deep image synthesis to generate artificial stimuli that distinctly engage encoding models of amygdala subregions that systematically differ from one another in terms of their low-level visual properties. These findings characterize how the amygdala compresses high-dimensional sensory inputs into low-dimensional representations relevant for behavior.

## Introduction

Animals navigate complex environments which contain diverse threats and opportunities for reward. Succeeding at this task depends on the amygdaloid complex—a subcortical cluster of nuclei in the medial temporal lobe (Swanson and Petrovich, 1998; Murray and Wise, 2004). The amygdala receives inputs from multiple sensory modalities (McDonald, 1998; Sah et al., 2003; Janak and Tye, 2015) and is a convergence zone with connections to much of cortex, subcortex, and midbrain systems involved in motivated behavior and autonomic control (Pessoa and Adolphs, 2010). The primates amygdala receives information about the environment predominantly from the ventral visual stream (Pessoa and Adolphs, 2010; Kravitz et al., 2013). Through computations performed on these and other inputs, the amygdala is thought to detect events of biological relevance and prepare animals to react appropriately (Sander et al., 2003; Cunningham and Brosch, 2012).

Human neuroimaging has shed light on amygdala function by examining its sensitivity to differences in reward, threat, valence, salience, and affective intensity. Typical experiments identify associations between different stimulus properties and amygdala responses. Meta-analytic summaries of this work show that the amygdala is sensitive to a wide array of biologically relevant inputs (Costafreda et al., 2008; Vytal and Hamann, 2010; Lindquist et al., 2012, 2016; Kragel and LaBar, 2016). One explanation of these findings is that the amygdala is involved in multiple functions, and that different neural ensembles process different stimulus properties relevant for distinct behaviors. However, identifying the set of variables that best explain amygdala function has been a challenge, as most studies only manipulate one or a few variables at a time, limiting strong inferences on amygdala specialization.

An alternative way to understand amygdala function is through systems identification. This approach involves building models of a system from measurements of its inputs and outputs. From this perspective, a complete understanding of amygdala function would comprise a model that transforms amygdala inputs (e.g., projections originating in the ventral visual stream) onto output variables conveyed to downstream structures (e.g., the hypothalamus, striatum, and midbrain structures). Compared to conventional approaches that involve manipulating a small number of variables and measuring changes in amygdala activity, systems identification requires experiments with complex sensory inputs that better match the diversity of amygdala inputs. The performance of computational models that predict amygdala responses to a given set of sensory inputs provides a metric for quantifying our understanding of brain function.

Here we probe multiple aspects of amygdala function from a systems identification perspective. Given evidence that the majority of sensory inputs to the primate amygdala originate from the ventral visual cortex (Kravitz et al., 2013), we predict that a computational proxy of the ventral stream should be sufficient to predict amygdala responses to emotionally evocative stimuli. Because sensory inputs predominantly project to the basal and lateral nuclei, whereas other nuclei are involved in different functions, prediction accuracy should systematically differ across amygdala subregions. We test these predictions using a combination of human neuroimaging, computational models of visual processing, and self-reported emotion. We analyze human brain responses to a full-length motion picture film (Aliko et al., 2020) and develop linear encoding models to predict amygdala responses using a deep convolutional neural network (Kragel et al., 2019) trained to recognize the emotional content of scenes.

We validate these models in two *in silico* experiments focused on prediction and control. First, we examine whether the models predict valence and arousal ratings in response to naturalistic images from two affective image databases (Bradley and Lang, 2007; Kurdi et al., 2017). Second, we use deep image synthesis (Nguyen et al., 2016; Bashivan et al., 2019) to generate visual stimuli that maximally engage amygdala subregions and subsequently identify which visual properties make them distinct. Collectively, these tests establish a framework for understanding amygdala function by characterizing how it transforms visual inputs into low- dimensional representations that can be used to guide behavior.

## Methods

### Development of Amygdala Encoding Models

We fit encoding models (Naselaris et al., 2011) to develop image computable models that take images presented to participants as inputs and predict amygdala responses (Figure 1). Based on anatomical and functional connectivity (Amaral and Price, 1984; Kravitz et al., 2013), we used a deep convolutional neural network that approximates the primate ventral visual stream (Kar et al., 2019) as it extracts highly processed visual features that are fed forward into lateral amygdala. We fit models using brain responses to naturalistic audiovisual stimuli with rich socioemotional content known to engage the amygdala.

**Figure 1.**
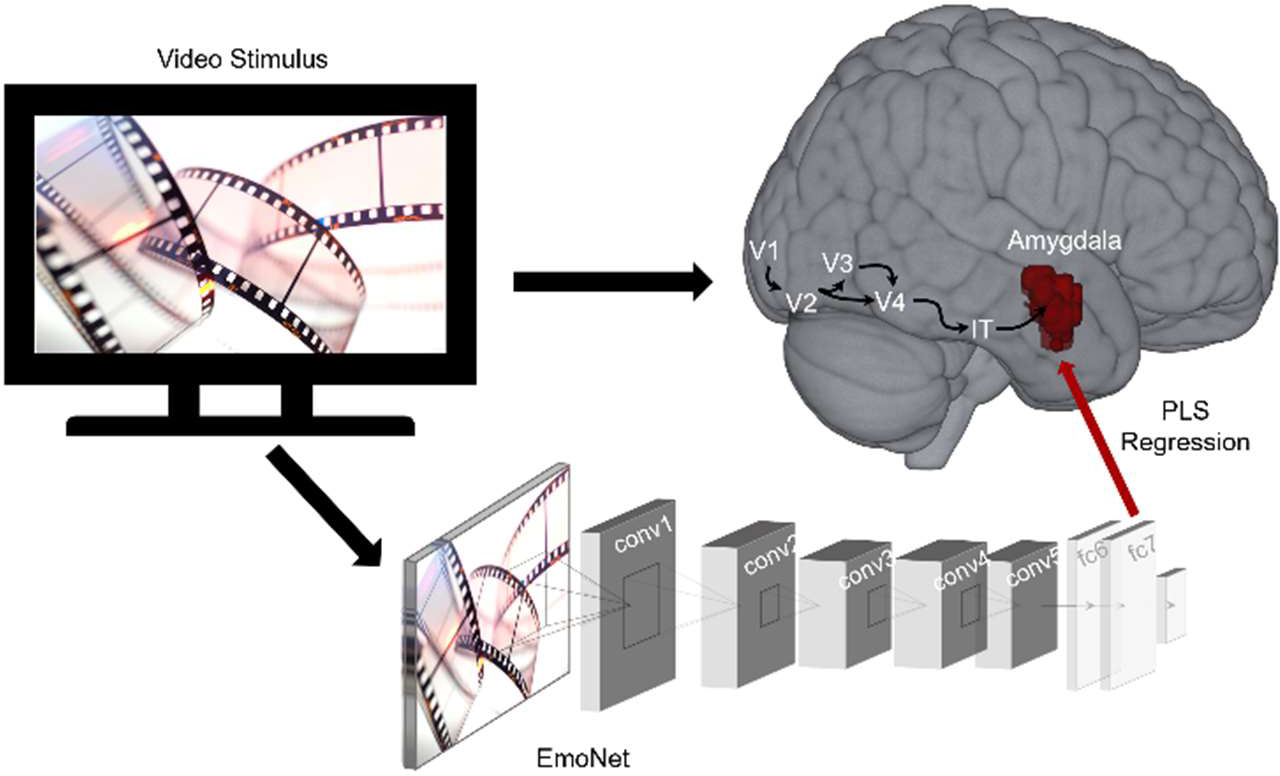
Schematic of encoding model workflow. A full-length movie was shown to participants concurrent with fMRI and was input to a deep convolutional neural network to extract features from frames of the video stimulus. Partial least squares regression identified a mapping between visual features and amygdala response patterns for each subject (*N* = 20). V1-V4: visual areas 1-4; IT: inferotemporal cortex; conv: convolutional layer; fc: fully connected layer; PLS: partial least squares.

### Neuroimaging Experiment

Functional magnetic resonance imaging (fMRI) data for this study were sampled from the Naturalistic Neuroimaging Database (NNDb) (Aliko et al., 2020). Detailed descriptions of the participants, the paradigm used for data acquisition, and the preprocessing of the fMRI data have been described elsewhere (Aliko et al., 2020; Soderberg et al., 2023). Briefly, blood oxygen level dependent (BOLD) data from 20 subjects viewing a full-length motion picture film *500 Days of Summer* was previously collected in a 1.5 T Siemens MAGNETOM Avanto with a 32 channel head coil (Siemens Healthcare, Erlangen, Germany) and consequently used for this study.

### Feature Extraction

We used a deep convolutional neural network, EmoNet (Kragel et al., 2019), as a feature extractor for encoding models. This model was finetuned from AlexNet (Krizhevsky et al., 2012) to classify emotional scenes and consists of five convolutional layers and three fully connected layers. We passed every fifth frame of the movie shown to participants during scanning as inputs to EmoNet and extracted features from the penultimate layer fc7 because this layer best approximates later stages of processing in the ventral visual pathway (Horikawa and Kamitani, 2017; Kragel et al., 2019).

### Regions of Interest

We modeled fMRI signal localized to amygdala masks based on cytoarchitecture (Amunts et al., 2005) and include the bilateral amygdala (247-252 voxels) and amygdala subregions (the basolateral complex (LB), the centromedial nucleus (CM), the superficial (SF) group, and the amygdalostriatal transition zone (AStr); 29 to 178 voxels). Some participants had partial coverage in some regions of interest (4 out of 20 subjects had < 252 voxels for the amygdala). We also fit encoding models for multiple control regions, including early cortical visual areas (V1-V3; 3,061-3,069 voxels) and the inferotemporal cortex (TE2, TF; 700-1010 voxels) examined bilaterally as delineated by multi-modal parcellation (Glasser et al., 2016).

### Model Specification

After extracting the image features from the movie, we convolved these features to account for the hemodynamic time delay of the BOLD data using a canonical double gamma response function (Friston, 2007). We specified separate partial least squares regressions (Wold et al., 2001) for each subject to obtain regression coefficients (beta estimates) for encoding models. We used the time-matched image features from the movie as the predictor variable and the observed BOLD activations masked by the voxels of the amygdala and other control regions of interest as the outcome variable. We specified encoding models for each region of interest (amygdala and its subregions, visual cortex, and inferotemporal cortex) for each subject that predict activations in each region of interest in response to the dynamic visual stimuli.

### Model Estimation

After specifying these encoding models, we used 5-fold cross validation to estimate the correlation between voxelwise encoding model predictions and the observed activations for each subject. Multivariate mappings were identified between visual features and BOLD response patterns using partial least squares regression. Regression models were regularized by retaining 20 components. We calculated the correlation between the predicted and observed activations for each voxel and normalized the coefficients using Fisher’s Z transformation for inference.

### Statistical Inference

To assess whether performance was above chance levels, we conducted one-sample *t*- tests on voxel-wise and region-average data. Voxel-wise inference was performed using false discovery rate correction with a threshold of *q* < .05. To test for differences in predictive performance across amygdala subregions, we performed a one-way repeated measures ANOVA. We specified planned contrasts that compared the performance of amygdala encoding models in the LB subregion with other amygdala subregions (CM, SF, AStr), the performance of the CM subregion to the SF and AStr subregions, and the performance in the SF subregion to the AStr subregion.

### Evaluating Encoding Model Responses to Affective Images

We validated encoding models using naturalistic images from standardized affective image databases (i.e., the International Affective Picture System (Bradley and Lang, 2007) and the Open Affective Standardized Image Set (Kurdi et al., 2017)). The goal of this experiment was to determine whether the predicted activations from our encoding models would behave similarly to human brains—exhibiting increased engagement along the dimensions of valence or arousal (Lindquist et al., 2016). Because it is well-established that differences in low-level visual properties are associated with alterations in valence and arousal in these databases (Anders et al., 2008; Styliadis et al., 2014; Bonnet et al., 2015; Hartling et al., 2021), we also accounted for variation with low-level visual features, namely color (red, green, blue) and spatial power (high and low spatial frequencies).

We used the naturalistic images as inputs to encoding models and tested for associations with normative valence and arousal ratings, and their interactions. We performed this analysis on both the IAPS and OASIS datasets. For each region, the responses to every image for each of the 20 encoding models (one per subject) were obtained by multiplying the activation produced in layer fc7 of EmoNet with the regression coefficients of that subject’s encoding model. We obtained the normative valence and arousal ratings for each of the naturalistic images. We then extracted the low-level visual features of color intensity (red, blue, and green) and spectral power (high and low frequencies). We produced color histograms for each IAPS and OASIS image and calculated the median value for each color. We calculated the power spectral density of each image using Fast Fourier Transform and then defined low frequencies as those with a radius < 30 pixels in Fourier space and high frequency as those with a radius > 50 pixels.

We conducted linear regression models with either the amygdala or visual cortex as the outcome variable using standardized predictor variables of valence ratings, arousal ratings, the interaction between valence and arousal (coded such that more positive and arousing images would produce the strongest response in an encoding model) and controlling for the low-level visual features of the median intensity of red, green, and blue, and the power in high and low spatial frequency bands. We used the fitlme function in MATLAB to build the models for each subject and performed group *t*-tests on the betas, treating subject as a random variable.

### Controlling Amygdala Encoding Model Responses using Deep Image Synthesis

After verifying the performance of our encoding models on naturalistic images, we wanted to synthesize artificial stimuli that could engage the encoding models of the amygdala and different amygdala subregions. Previous studies have demonstrated related approaches can target activation to specified units within the visual cortex in both humans and non-human primates (Nguyen et al., 2016; Bashivan et al., 2019; Xiao and Kreiman, 2020; Wang and Ponce, 2022). Here we extended this method to generate artificial stimuli that would target the amygdala (Figure 2). We used a deep generator network trained on ImageNet (Nguyen et al., 2016) and the outputs of our amygdala encoding models to map activation in layer fc7 of EmoNet as the objective for activation maximization. This was accomplished by computing the dot product with different sets of encoding model regression coefficients (beta estimates) that predicted the responses of different amygdala voxels. Optimization was performed using an evolutionary algorithm (Wang and Ponce, 2022) implemented in Python (https://github.com/Animadversio/ActMax-Optimizer-Dev). We used this procedure to generate artificial stimuli targeting the average amygdala response, individual amygdala subregions (LB, CM, SF and AStr), visual cortex, and inferotemporal cortex.

**Figure 2.**
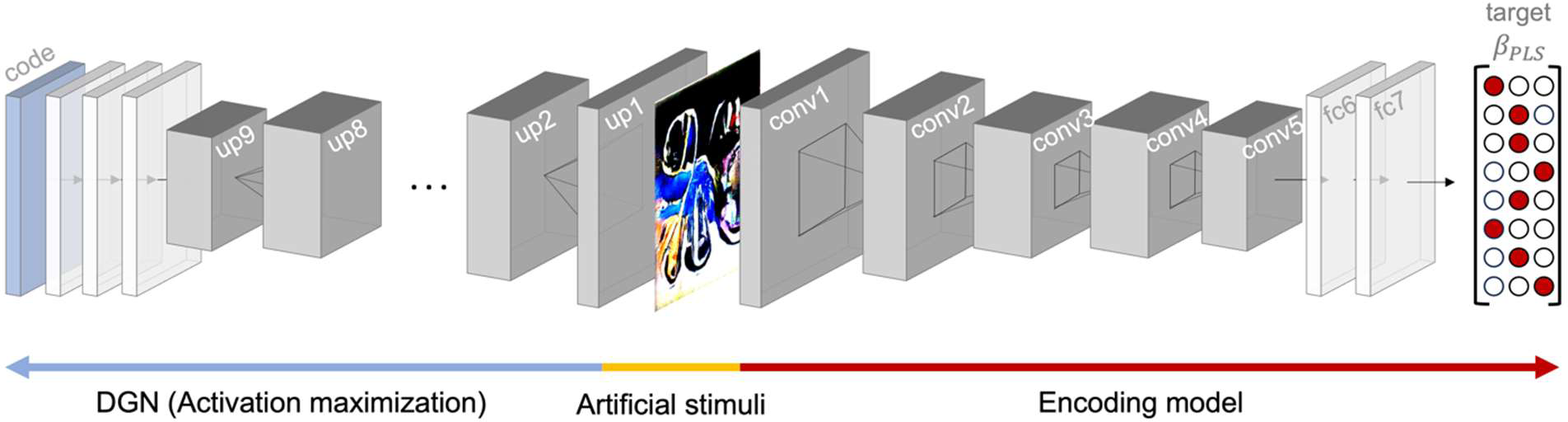
Artificial image synthesis procedure. A deep generator network (DGN; blue arrow) initialized with a random code produces an artificial stimulus (yellow) that is fed as input into the encoding model (red). Beta estimates specifying the relationship between unit activity in the deep convolutional network and BOLD response patterns serve as the target for activation maximization. Forward and back propagation update the code to modify and generate an artificial stimulus that maximizes activation patterns in the target region. up: upconvolutional layer; conv: convolutional layer; fc: fully connected layer.

Artificial stimuli were generated with a random starting seed for each image. The optimization algorithm did not converge for some seeds (producing an identical image); these images were excluded from subsequent analyses. As a result, 4-5 different artificial stimuli were generated for each region of interest for each subject, resulting in 80 artificial stimuli synthesized per region of interest. An exception to this was the artificial stimuli generated for the inferotemporal cortex; because it was used as a control region, 9 artificial stimuli were generated for each subject resulting in a total of 160 artificial stimuli.

To assess the selectivity of encoding models, we assessed whether they responded differentially to generated stimuli optimized for different regions of interest. Following the same procedures used evaluate the naturalistic stimuli, we fed the artificial stimuli (n = 686) into all encoding models and obtained a predicted activation for each of the artificial stimuli. We also characterized low-level visual features such as color (red, blue, and green) and spectral power (high and low frequencies) found in the synthesized artificial stimuli as predictors in our models. We performed linear regressions on standardized variables to confirm that the synthesized images activated their intended targets. We fit mixed-effects models for each subject with target region for image synthesis (on vs off target), the subject used for image synthesis, and the low-level visual features described above as predictors for within subject fixed effects. Separate models were run to predict the activation of the amygdala, each of its subregions (LB, CM, SF and AStr), and visual cortex. We used the fitlme function in MATLAB to build each model and compared the betas of the models using *t*-tests.

To evaluate the discriminability of artificial stimuli, we performed a supervised classification and examined confusions between the predicted and actual region targeted for optimization. Multi-way classification models were estimated using partial least squares discriminant analyses (7 components). Generalization performance was estimated using 5-fold cross validation. Confusions between different image classes were assessing using a hierarchical approach in a 7-way classification, with the number of clusters set to be the maximum number of clusters in which all pairs of clusters are statistically discriminable from one another. To visualize the results of this analysis, we generated a *t*-SNE plot (Maaten and Hinton, 2008) based on the model predictions for each of the artificial stimuli.

## Results

We found that visual features captured by deep convolutional neural networks are encoded in amygdala responses to naturalistic, dynamic videos. Voxel-wise tests showed that the mean performance of encoding models was well above chance (Figure 3). A mixed effects model revealed that predictions of the average amygdala response were also above chance (*β̂* = .049, *SE* = .0053, *t(53)* = 9.27, *p* < .001), and that there were marked differences in performance across amygdala subregions (ΔBIC = 23.5, Likelihood Ratio = 36.5, *p* < .001). The first contrast comparing LB to the other three subregions did not result in statistical significance (*β̂* = -.0012, *SE* = .0012, *t(53)* = -1.04, *p* = .304). The other two contrasts indicated differences between the performances of CM and the average of SF and AStr (*β̂* = .0036, *SE* = .0015, *t(53)* = 2.39, *p* = .020), and between the SF and AStr (*β̂* = .017, *SE* = .0026, *t(53)* = 6.47, *p* < .001). Post-hoc tests indicated that there were differences between CM and AStr (*β̂* = .027, *SE* = .0050, *z* = 5.45, *p* < .001), SF and AStr (*β̂* = .033, *SE* = .0050, *z* = 6.64, *p* < .001), SF and LB (*β̂* = .018, *SE* = .0054, *z* = 3.33, *p* = .005), LB and AStr (*β̂* = .015, *SE* = .0054, *z* = 2.84, *p* = .023), but not between CM and LB or between SF and CM. Thus, the sets of voxels for SF and CM exhibited the highest performance, followed by voxels for LB, and then the voxels for AStr.

**Figure 3.**
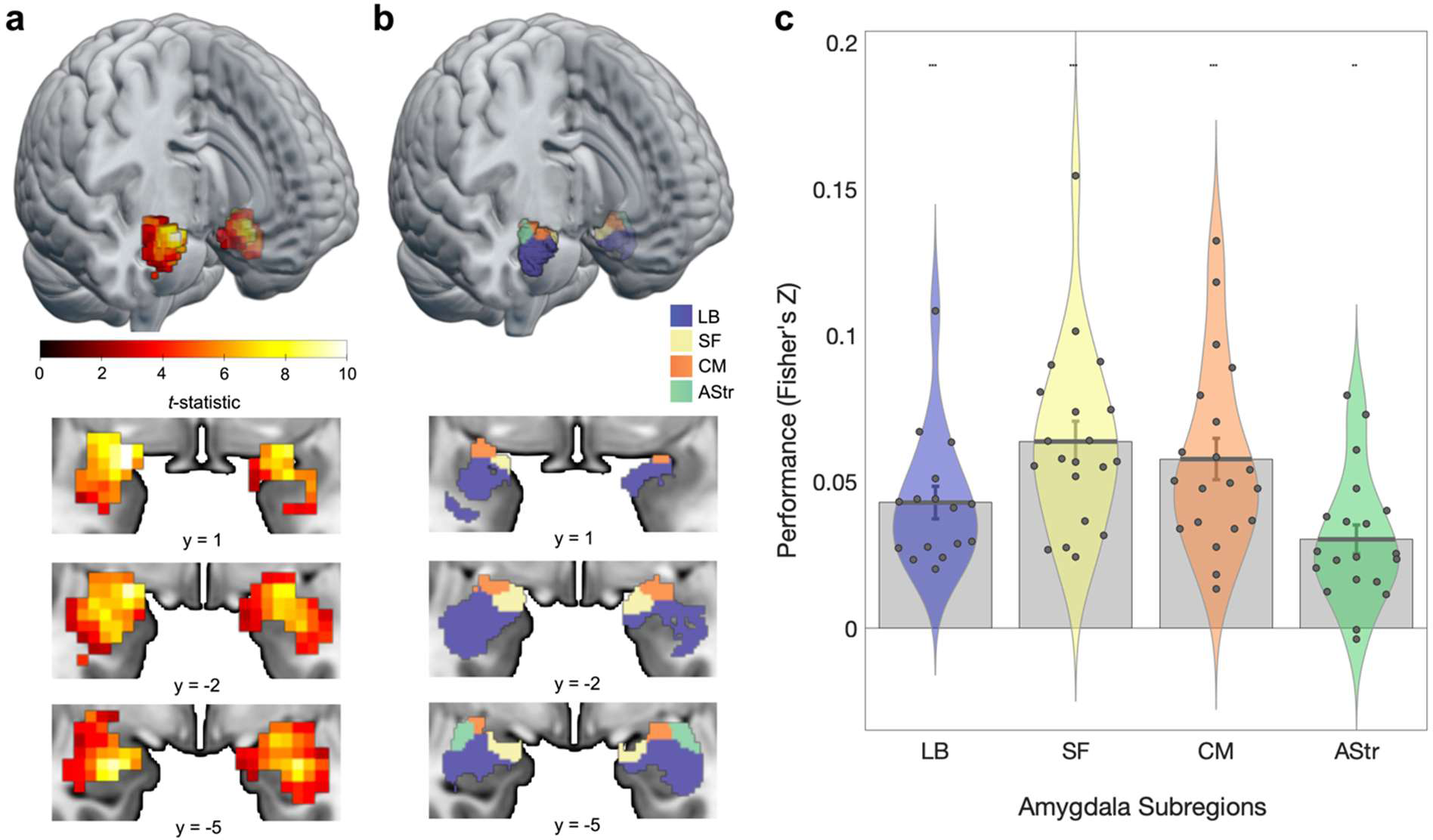
**ANN-based encoding models predict human amygdala responses to naturalistic videos**. **a**) Amygdala activation is predicted by encoding models fit on naturalistic videos (group *t*-statistic computed on the cross-validated correlation between predicted and observed BOLD responses). Maps are displayed with a threshold of *q*FDR < .05. **b**) Rendering of amygdala parcellation (Julich-Brain Cytoarchitectonic Atlas). Blue, LB: laterobasal; yellow, SF: superficial; Orange, CM: centromedial; green, AStr: amygdalostriatal. (**c**) Violin plots of average predictive performance of encoding models in each subregion. Each point corresponds to a single subject (*N* = 20). Error bars reflect standard error of the mean. \**p* < .05, ** *p* < .01, *** *q*FDR < .05

### Predicting the response of amygdala-based models along dimensions of valence and arousal

We validated our encoding models on images from the IAPS and OASIS datasets that have been shown to produce increases in amygdala activity (Britton et al., 2006; Haj-Ali et al., 2020; Hartling et al., 2021) along the dimensions of valence (Garavan et al., 2001; Anders et al., 2004, 2008; Mather et al., 2004; Aldhafeeri et al., 2012; Styliadis et al., 2014) and arousal in humans (Canli et al., 2000; Kensinger and Schacter, 2006). Consistent with previous fMRI studies that show increased amygdala responses to positively valent stimuli, we found that the amygdala encoding model captured linear increases in valence (*β̂* = .0095, *t(19)* = 3.13, *p* = .006, *d* = 0.70; Figure 4). Encoding model responses did not track arousal (*β̂* = .0006, *t(19)* = 0.18, *p* = .861, *d* = 0.04) or the interaction between valence and arousal (*β̂* = -.0034, *t(19)* = -1.48, *p* = .155, *d* = -0.33). Moreover, we found that the amount of red color (*β̂* = .0073, *t(19)* = 2.24, *p* = .037, *d* = 0.50) and high frequency spatial power (*β̂* = .0211, *t(19)* = 2.96, *p* = .008, *d* = 0.66) within images also predicted activations in amygdala models.

**Figure 4.**
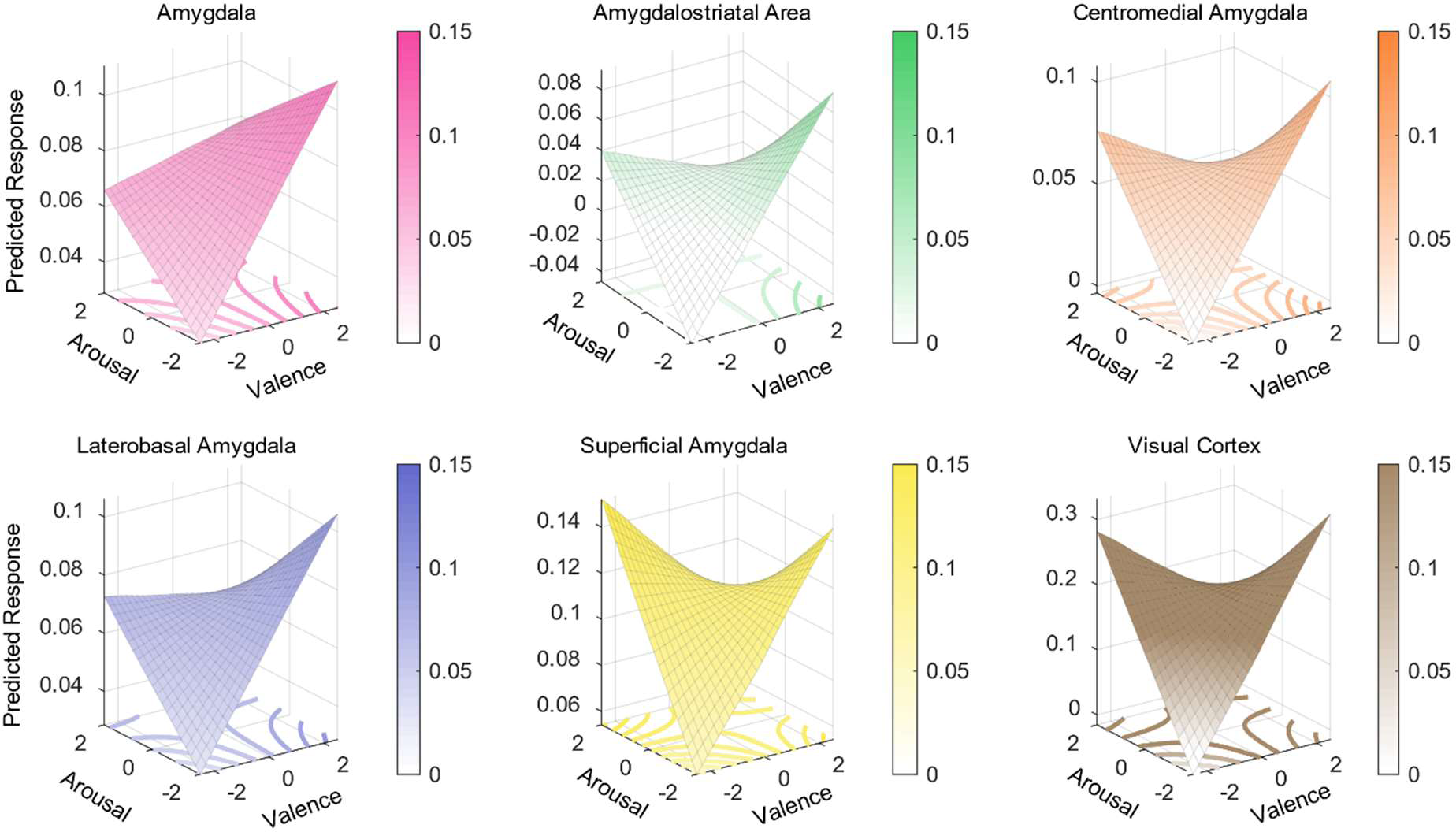
Amygdala encoding model responses to images from the standardized affective images. The predicted response to images from the International Affective Picture System (IAPS) and the Open Affective Standardized Image Set (OASIS). Predictions were generated from regression models predicting encoding model responses based on valence, arousal, the interaction between valence and arousal. Surface plots show responses averaged across the entire amygdala, visual cortex, and within amygdala subregions.

Given recent findings from multivariate decoding studies demonstrating that the amygdala encodes valence along a single dimension that ranges from unpleasantness to pleasantness (Jin et al., 2015; Tiedemann et al., 2020), we performed a series of regressions examining associations with valence separately for negative (z < 0), neutral (absolute value of z < 1), and positive (z > 1) images. If the amygdala encoding model predicts valence across the full valence spectrum using a single continuous representation, then we would expect all three regressions to exhibit a positive relationship. Alternatively, the amygdala may encode coarse-grained differences in valence extremes using a discontinuous function, consistent with bivalent models of affect (Bradburn, 1969; Watson and Tellegen, 1985; Cacioppo et al., 2012; Mattek et al., 2017).

Consistent with the latter hypothesis, we found amygdala encoding models respond to valence in a piecewise, discontinuous manner. Increasingly negative images produced greater activations in the encoding model (*β̂* = -.0140, *t(19)* = -2.59, *p* = .018, *d* = -0.58). Valence coding shifted within the neutral range, as more positive images produced greater activations (*β̂* = .0187, *t(19)* = 4.37, *p* < .001, *d* = 0.98). This coding continued for more extreme positive images, as they produced greater activations in the encoding model (*β̂* = .0126, *t(19)* = 2.46, *p* = .024, *d* = 0.55). These results suggest that the encoding model captures coarse-grained differences between valence extremes and also a more fine-grained, nonlinear representation of valence.

As our overarching hypothesis is that the amygdala functions to select among many possible behaviorally relevant sensory features, we next examined whether affective variables encoded in the activity of the visual cortex differed from those of amygdala responses.

Examining relationships between visual cortex encoding model predictions and normative affective variables, we found a positive association with valence (*β̂* = .0188, *t(19)* = 5.06, *p* < .001, *d* = 1.13) and arousal (*β̂* = .0104, *t(19)* = 2.74, *p* = .013, *d* = 0.61), and a significant interaction (*β̂* = -.025, *t(19)* = -8.05, *p* < .001, *d* = -1.80), such that the encoding model responded more with increasing arousal for negative compared to positive stimuli. These results are broadly consistent with data showing that amygdala feedback modulates early visual responses (Liu et al., 2022) and that visual cortex encodes representations of multiple affective variables (Miskovic and Anderson, 2018; Kragel et al., 2019; Li et al., 2019; Bo et al., 2021).

To evaluate whether amygdala and visual cortex encoding of affective variables differed, we compared the strength of associations between regions. The amygdala encoding models had weaker associations with both valence (*β̂* = -.009, *t(19)* = -2.19, *p* = .041, *d* = -0.49) and arousal (*β̂* = -.010, *t(19)* = -2.25, *p* = .036, *d* = -0.50) compared to visual cortex models. Similarly, the amygdala models exhibited a weaker (less negative) interaction between valence and arousal compared to the visual cortex encoding models (*β̂* = .0218, *t(19)* = 6.10, *p* < .001, *d* = 1.36).

Given the functional heterogeneity of the amygdala and past evidence demonstrating interactions between valence and arousal (Winston et al., 2005), we next tested whether there were differences in the encoding of valence and its interaction with arousal in amygdala subregions. To this end, we fit separate encoding models for each amygdala subregion. We performed ANOVAs comparing activations between subregions and found that responses related to valence did not differ across subregions (*F(1,19)* = 3.95, *p* = .062), whereas the interaction between valence and arousal varied across subregions (*F(1,19)* = 7.45, *p* = .013). Exploratory post-hoc tests did not reveal any significant effects after correcting for multiple comparisons, although AStr and LB demonstrated a difference with a modest effect size (*β̂* = 0.0045, *SE* = .0023, *p* = .254, 95% CI = [-.0021, .0111], *d* = -.429; Table 1).

**Table 1.** Effects of valence and arousal on amygdala subregions. LB: laterobasal; SF: superficial; CM: centromedial; AStr: amygdalostriatal.

| Subregion | Valence (main effect) |  |  |  | Valence by Arousal (interaction) |  |  |  |
| --- | --- | --- | --- | --- | --- | --- | --- | --- |
|  | Coefficient | SE | p | Cohen's d | Coefficient | SE | p | Cohen's d |
| LB | 0.0068 | .0027 | .022 | .56 | -0.0043 | .0017 | .021 | -.56 |
| SF | 0.0030 | .0078 | .709 | .08 | -0.0077 | .0063 | .237 | -.27 |
| CM | 0.0073 | .0037 | .064 | .44 | -0.0075 | .0039 | .071 | -.43 |
| AStr | 0.0102 | .0035 | .009 | .65 | -0.0088 | .0023 | .001 | -.85 |

### Controlling encoding models of distinct amygdala subregions

To further evaluate regional specificity, we generated artificial stimuli optimized to activate anatomically defined amygdala subregions (i.e., LB, SF, AStr, and CM amygdala; Figure 5). We then compared the activity produced by on- vs. off-target artificial stimuli within the respective encoding models. This analysis revealed that artificial stimuli selectively engaged on-target subregions compared to off-target subregions (AStr: *β̂* = .026, *t(19)* = 4.63, *p* < .001, *d* = 1.04; CM: *β̂* = .031, *t(19)* = 5.89, *p* < .001, *d* = 1.32; LB: *β̂* = .009, *t(19)* = 2.22, *p* = .039, *d* = 0.50), with the exception of SF (*β̂* = .023, *t(19)* = 1.31, *p* = .205, *d* = 0.29). A supervised classification analysis revealed all image types were distinct from one another in pairwise comparisons, with the exception of the artificial stimuli generated to target the LB and SF subregions. The six distinct image clusters could be discriminated from one another in a 6-way classification with 71.7 ± 1.7% (*SE*) accuracy (chance accuracy = 21.96 ± 16.4%), demonstrating a high degree of functional specialization (Figure 6).

**Figure 5.**
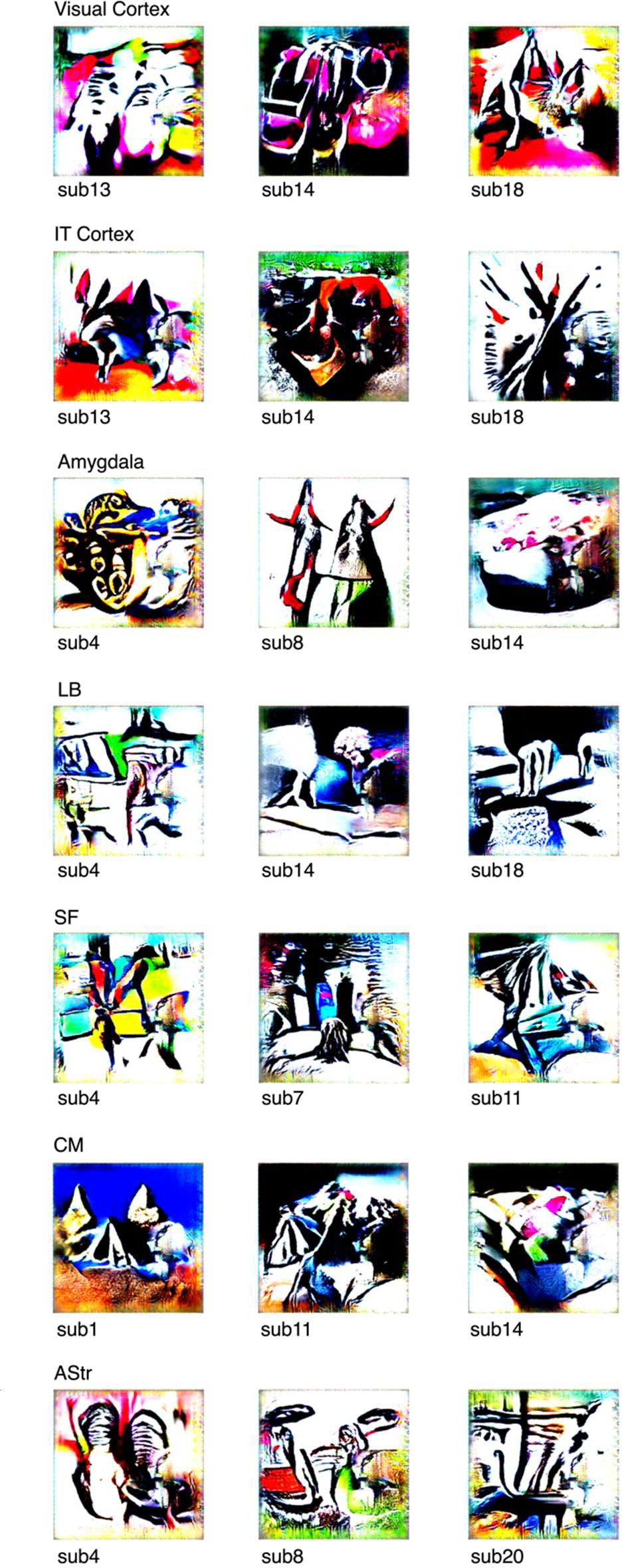
Representative artificial stimuli for each target region. LB: laterobasal; SF: superficial; CM: centromedial; AStr: amygdalostriatal; sub: subject.

**Figure 6.**
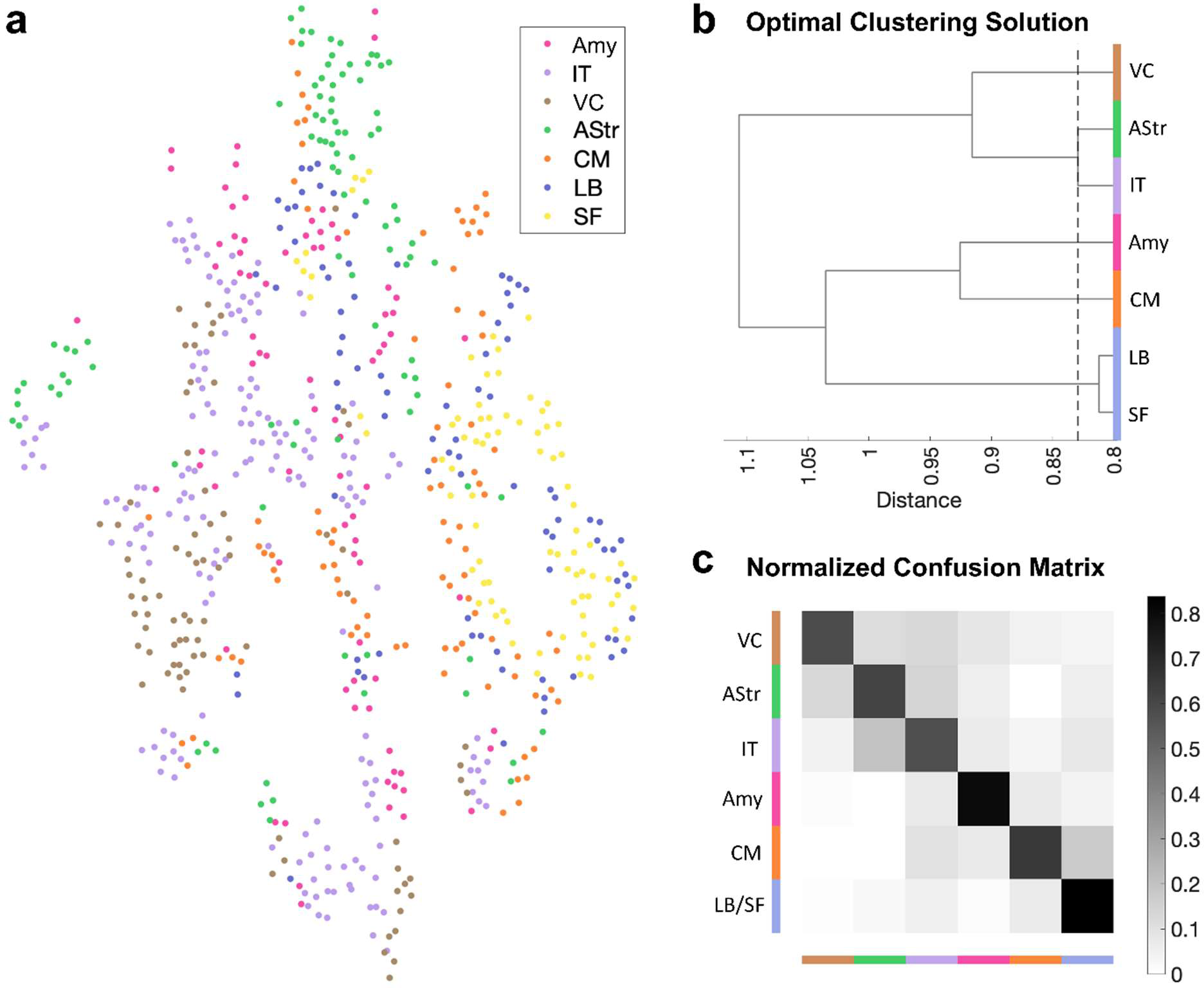
**ANN-generated stimuli selectively engage encoding models of different regions of interest. a**) *t*-SNE plot, **b**) optimal clustering solution, and **c**) normalized confusion matrix of predicted activations of stimuli in encoding models color-coded by region of interest. Confusion matrix shows above chance performance. amy: whole amygdala; IT: inferotemporal cortex; VC: visual cortex; AStr: amygdalostriatal transition zone; CM: centromedial amygdala; LB: laterobasal amygdala; SF: superficial amygdala.

## Discussion

We found that amygdala processing can be characterized using a systems identification framework. Encoding models using features from deep convolutional neural predicted BOLD activity within multiple amygdala nuclei during free viewing of a cinematic film. In independent validation tests, the amygdala encoding model consistently responded to differences in valence and its interaction with arousal, the amount of red color, and high spatial frequency power of affective images, consistent with prior work investigating amygdala responses to these stimuli (Garavan et al., 2001; Anders et al., 2004, 2008; Styliadis et al., 2014). Furthermore, stimuli synthesized to engage amygdala subregions were visually distinct, alluding to differences in the specialization of amygdala subregions. We take these findings to show that one function of the amygdala is to transform sensory inputs from the ventral visual stream to produce representations related to valence.

Our findings demonstrate how encoding models can be used to characterize the interface between sensory pathways and downstream regions involved in cognition and emotion. A large body of work has used hand-engineered (Jones and Palmer, 1987; Lee, 1996; Dumoulin and Wandell, 2008) and data-driven (Fukushima, 1988; Riesenhuber and Poggio, 1999) features to characterize the primate visual system. Deep convolutional neural networks have been developed as models of the ventral visual stream—providing a better match to the complexity of biological systems underlying perception (Yamins and DiCarlo, 2016; Kar et al., 2019). The existing literature work has generally focused on identifying the best one-to-one mappings between specific features and the responses of distinct visual areas to carefully controlled stimuli, with the goal of identifying a fully mappable model of the visual system (Yamins and DiCarlo, 2016) ranging from the retina to the anterior temporal lobe. Here we explored mappings that diverge from ventral stream involvement in visual recognition to characterize a system central to emotional behavior, the amygdaloid complex (O’Neill et al., 2018).

Characterizing amygdala function using an encoding model framework is a departure from common methods that involve measuring amygdala responses to one or a few variables at a time (Garavan et al., 2001; Anderson et al., 2003; Anders et al., 2004, 2008; Kensinger and Schacter, 2006; Styliadis et al., 2014; Jin et al., 2015; Haj-Ali et al., 2020; Tiedemann et al., 2020). Whereas conventional studies are built upon well-founded assumptions that the amygdala is involved in processing specific variables such as threat, reward, pleasure, and intensity, among others, we relaxed these constraints and predicted that amygdala responses can be approximated as an image-computable function of signals present in the sensory array. Thus, although we did not assume any specific variable was encoded in amygdala activity, we found that amygdala encoding models were sensitive to variation in the normative valence and arousal evoked by images.

In line with our observation that the average response of the amygdala encoding model increased from negative to positive extremes of the valence continuum, recent multivariate decoding studies have shown that the amygdala unidimensionally represents the valence of odors (Jin et al., 2015) and images of food (Tiedemann et al., 2020). Together, these findings are broadly consistent with studies reporting the amygdala is involved in reward learning and evaluating social images (Baxter and Murray, 2002; Adolphs and Spezio, 2006). They are also congruent with work in nonhuman primates showing that both pleasant and unpleasant stimuli engage distributed neural populations in the amygdala (Paton et al., 2006; Belova et al., 2008), and with fMRI evidence showing that the amygdala participates in a distributed network of brain regions sensitive to fluctuations in hedonic valence (Kragel et al., 2023).

In addition to variation related to valence extremes, we observed nonlinearities in encoding model responses to affective images, such that responses were greater for highly valent compared to neutral stimuli. This pattern of results has been observed in response to olfactory (Winston et al., 2005) and auditory (Fecteau et al., 2007) stimulation. Whereas unidimensional coding of valence was widespread throughout the amygdala, we found this interactive effect modestly differed across amygdala subregions, with the largest effect in the amygdalostriatal transition area, a region that encodes the valence of threatening stimuli and is important for the expression of conditioned defensive behavior in nonhuman animal models (Goto et al., 2022; Mills et al., 2022). It is possible that overlapping neural populations in the amygdala relate to valence in different ways, based on contextual factors that influence connectivity with distributed brain networks (Gothard, 2020). For instance, one recent study (Čeko et al., 2022) identified representations of negative affect from different sensory origins (visually evoked and domain-general across somatic, thermal, visual, and auditory sources) and non-specific arousal that were distributed across brain systems, yet overlapped in the amygdala. The amygdala activity captured by our encoding models could reflect visual-specific or domain-general coding of affect; adjudicating between these alternatives requires further study.

We found that stimuli generated to selectively engage amygdala subregions were clustered such that stimuli generated to engage the input centers of the amygdala (such as the LB) were distinct from output centers of the amygdala (such as the CM and AStr). This result is broadly consistent with models of amygdala processing that suggest the amygdala identifies a subsets of sensory variables that are relevant for learning and motivating behavior (Pessoa, 2010; Sladky et al., 2024). However, the overall distinctiveness of synthetic stimuli raises other possibilities. Differences in synthetic stimuli could result from local processing within the amygdala or connections to the amygdala bypass the laterobasal complex and directly influence population activity.

Despite exhibiting large effect sizes, voxel-wise predictions were far from explaining all amygdala activity. This is perhaps unsurprising, given the complexity of the movie stimulus and the relative simplicity of the encoding model used. As we developed encoding models using static visual features useful for classifying emotional scenes, amygdala responses to emotional stimuli from other sensory modalities (e.g., auditory and linguistic signals), those that habituated over time, or were dependent on learning taking place over the course of the movie stimulus could not be predicted using our approach. We anticipate that amygdala responses influenced by these factors can be characterized using similar approaches, given connections between the amygdala and brain regions involved in reinforcement learning, audition, and language (Price, 2003; Koelsch et al., 2013; Abivardi and Bach, 2017), and the success of computational models in characterizing the function of these systems (Yamins and DiCarlo, 2016; Cross et al., 2021).

Amygdala encoding models were trained on the visual input of one full-length motion picture film, *500 Days of Summer*, and on the corresponding brain data of 20 subjects viewing this movie. This full-length movie is sufficiently complex with both positive and negative valence scenes, faces, and other visual content, although it may have been limited in its ability to evoke robust and varied emotional experiences, including acute fear (Hudson et al., 2020).

Future studies using different movies, videos, or other dynamic visual stimuli to train encoding models are needed to identify the set of variables encoded by the amygdala, and to assess the extent to which they are context dependent or generalize across stimulus types (Čeko et al., 2022) and situations (Kragel et al., 2023).

In conclusion, our study shows that the amygdala encodes multiple features of visual stimuli, ranging from low-level features such as color and spectral power to more complex features along the dimension of valence, with marked differences between the features that individual amygdala subregions represent. Thus, perhaps what is driving the amygdala can be thought of as something beyond a single dimension or a handful of constructs, but rather a large array of features yet to be identified and objectively examined to understand how the amygdala coordinates emotional behavior.

## Author Contributions

Conceptualization, methodology, formal analysis, writing, validation, and visualization, PAK; conceptualization, formal analysis, writing, and visualization, GJ.

## Data Availability

The fMRI data used to fit encoding models is available at https://openneuro.org/datasets/ds002837/versions/2.0.0. Data used for fine-tuning EmoNet are available upon request from https://goo.gl/forms/XErJw9sBeyuOyp5Q2. Other data relevant to this project is available at https://osf.io/r48gc/.

## Code Availability

Code for all analyses will be made available upon publication at GitHub at https://github.com/ecco-laboratory/AMOD. The code used for implementing EmoNet in Python is available at https://github.com/ecco-laboratory/emonet-pytorch.

## Conflict of interest statement

The authors have no competing interests to declare.

## Acknowledgements

This project was partially supported by grant R01MH134972 to PK. GJ was supported by grant T32NS096050.

## Notes

### Competing Interest Statement

The authors have declared no competing interest.

